# Connectome-wide association analysis identified significant brain functional alteration in adults with attention-deficit/hyperactivity disorder

**DOI:** 10.1101/2021.12.14.472533

**Authors:** Lu Liu, Di Chen, Fang Huang, Tianye Jia, Meirong Pan, Mengjie Zhao, Xuan Bu, Yufeng Wang, Miao Cao, Qiujin Qian, Jianfeng Feng

## Abstract

Adults with attention-deficit/hyperactivity disorder (ADHD), as an extreme-phenotype of ADHD, is still facing problems of inconsistency and undeciphered mechanisms for its neuropathology. To address this matter, our present study performed connecotome-wide voxel-based analyses with the resting-state fMRI data of 84 adults with ADHD and 89 healthy controls. We found that functional connectivity patterns of the left precuneus and the left middle temporal significantly altered in ADHD populations serving as potential neural biomarkers to distinguish ADHD with healthy controls, with subsequent seed-based analysis revealing the dysfunction of functional connections both intra- and inter-default mode and attention networks, among which middle temporal gyrus plays the key role of ‘bridge’ linking the default mode and attention networks. After cognitive behavioral therapy, two of these ADHD-altered functional connections ameliorated accompanied with improvement of ADHD core symptoms. Additionally, imaging genetic analyses also revealed close relationships between the observed brain functional alterations and ADHD-risk genes. Taken together, our findings suggested the interference of default mode on attention networks in adults with ADHD, which would be severing as a potential biomarker for both ADHD pathogenesis and treatment effects.

## Introduction

Attention-deficit/hyperactivity disorder (ADHD) was one of the common neurodevelopmental disorders. According to the current diagnostic criteria, ADHD should be of childhood onset (before the age of 12 according to the DSM-5) (American Psychiatric Association, 2013). Approximately ~50% of children with ADHD will persist into adulthood (Lara et al., 2009), while the worldwide prevalence of ADHD in adults at about 2.5% (Dobrosavljevic et al., 2020). Adults with ADHD could be considered to be a specific and refined subgroup with poor outcome. ADHD persist into adulthood showed stronger family aggregation than those remitted prior to adulthood (Chen et al., 2016). Hence, adults with ADHD, as the most severe form (Franke et al., 2012), could be considered to be as an extreme-phenotype of this disorder. Exploration of the underlying pathogenesis of this extreme-phenotype would significantly promote our understanding of the natural history of ADHD.

Brain functional alterations have been consistently proposed to be involved in the neurobiological underpinnings of ADHD. Compared with healthy controls, adults with ADHD indicated alteration of intra- and/or inter-functional connectivity in default mode network (DMN), dorsal attention network (DAN), ventral attention network (VAN) and affective network (Mccarthy et al., 2013; Gao et al., 2019; Cortese et al., 2020). In spite of effortful seeking, no specific key brain region has been confirmed in adults with ADHD, that a recent meta-analysis yielded negative findings (Cortese et al., 2020). Multiple factors might contribute the inconsistency among studies. One noteworthy methodological issue is that most studies were depend on a priori hypotheses, with specific brain regions based on empirical evidence as regions of interests (ROIs) or targeted voxels. Whereas, ADHD, as a complex disorder, should be related with abnormalities in large-scale neural networks and concurrent activity across multiple distributed brain networks. Correspondingly, connectome-wide association studies (CWAS) with hypothesis-free would potentially help us to excavate some hidden brain signals related with ADHD from the whole-brain scale. Multivariate distance matrix regression (MDMR; Shehzad et al., 2014), an effective multivariate approach, has been proved to be efficient for CWAS to identify phenotypic associations in the whole-brain connectome for psychiatric disorders (Brady et al., 2019), including ADHD (Shehzad et al., 2014; Yang et al., 2016). In our present study, we anticipated that using MDMR for a whole-brain voxel-based analysis would get some potential voxel-based biomarkers for adults with ADHD.

In addition to the seeking biomarkers of psychopathology by hunting of the state-related (ADHD versus healthy controls) brain functional features, the amelioration of these neural indicators by clinical intervention (medication or non-medicine treatment) for the disorder would promote the elucidation of neural mechanisms from the perspective of predictive validity. Indeed, the evidence from the existing literature have indicated that ADHD-related brain functional interactions could be modulated after medication (atomoxitine, methylphenidate or amphetamine) (Pereira-Sanchez et al., 2020). For non-medicine treatment, cognitive behavioral therapy (CBT), an effective treatment for adults with ADHD, could also ameliorate the intrinsic functional brain networks in adults with ADHD (Wang et al., 2016). In addition, ADHD is a complex disorder with higher heterogeneity. Then, genetic factors would definitely play critical role in the underlying mechanisms and should be considered as key determinants (Nymberg et al., 2013). Linking the observed phenotype-related brain abnormalities with genetic risks would further validate the observed brain functional alteration from the perspective of construct validity.

In our present study, first, we performed whole-brain voxel-based analysis using Multivariate Distance Matrix Regression (MDMR) in 84 adults with ADHD and 89 healthy controls to detect ADHD-altered resting-state functional connectivities (FCs) and identify the potential neural biomarker for adults with ADHD. Second, using a sub-sample from our previous CBT study (14 adults with ADHD, 25 healthy controls), we explored whether those altered brain functional features could be ameliorated by CBT from the perspective of predictive validity, to built a potential ‘brain-treatment’ relationship. Finally, the explorative relationship of these brain functional biomarkers with ADHD-risk genes (*MAOA, MAOB*) were examined to demonstrate the genetic substrates of brain functional features, which would promote the understanding the chain of ADHD’s pathogenesis more comprehensively.

## Methods and Materials

### Study Participants

The study samples were recruited from clinics in Peking University Sixth Hospital/Institute of Mental Health. 84 adults with ADHD and 89 adult healthy controls (HC) were included for the baseline study. All ADHD subjects were interviewed by experienced psychiatrists according to the criteria of the Diagnostic and Statistical Manual of Mental Disorders (4th ed.; DSM-IV; American Psychiatric Association [APA], 1994) using the Conners’ Adult ADHD Diagnostic Interview. In addition, structured clinical interview for DSM-IV Axis-I disorders (SCID-I), and Axis-II disorders (SCID-II) were conducted to assess comorbid disorders. All included participants were: (1) aged 18-45 years; (2) right hand dominant; (3) no history of severe physical disease; (4) a full-scaled IQ intelligence quotient (FSIQ) evaluated using the Wechsler Adult Intelligence Scale-Third Edition above 80. Schizophrenia, clinically significant panic disorder, severe major depression or pervasive developmental disorders were excluded. In addition, any history of psychiatric disorders were also excluded for HC. Among those subjects recruited for baseline study, 14 ADHD and 25 HC, who have participated in the CBT study (NCT02062411; Huang et al., 2019) and were with available fMRI data at both baseline and post-treatment, were included in follow-up study to test the predictive validity of the observed altered brain functional features. Self-reports of ADHD Rating Scale-IV (ADHD RS-IV) were used to assess the severity of inattentive symptoms, hyperactive/impulsive symptoms and total ADHD symptoms.

The demographic and clinical characteristics of both baseline and follow-up studies were summarized in **Table** 1. The mean frame-wise displacement (FD) of each participant was smaller than 0.5. The effects of mean FD, age, FSIQ, gender and education years were excluded as the covariates in the following analysis. This study was approved by the Ethics Committee of Peking University Sixth Hospital/Institute of Mental Health. Before participation in the study, all participants provided written informed consent.

**Table 1.** Demographic and clinical characteristics of baseline and follow-up studies.

|  | Baseline study |  |  |  | Follow-Up study |  |  |  |
| --- | --- | --- | --- | --- | --- | --- | --- | --- |
| | ADHD (n=84) | HC (n=89) | $\chi^2/t$ | P-value | ADHD (n=14) | HC (n=25) | $\chi^2/t$ | P-value |
| Male/Female | 62/22 | 66/23 | <0.01 | 0.958 | 8/6 | 14/11 | <0.01 | 0.945 |
| Age (Mean ± SD) | 25.80 ± 3.93 | 26.04 ± 3.63 | -0.43 | 0.668 | 25.29 ± 3.99 | 26.08 ± 3.07 | -0.70 | 0.491 |
| Education year (Mean ± SD) | 15.88 ± 2.11 | 17.20 ± 1.97 | -4.26 | <0.001 | 16.36 ± 2.68 | 17.68 ± 2.21 | -1.66 | 0.110 |
| FSIQ (Mean ± SD) | 120.94 ± 7.48 | 121.83 ± 7.50 | -0.78 | 0.435 | 120.85 ± 7.85 | 120.44 ± 7.10 | 0.17 | 0.870 |
| **ADHD symptoms (Mean ± SD)** |  |  |  |  |  |  |  |  |
| Inattentive | 17.70 ± 3.94 |  |  |  | 13.36 ± 3.82 |  |  |  |
| Hyperactive/Impulsive | 10.16 ± 5.11 |  |  |  | 9.14 ± 5.05 |  |  |  |
| Total | 28.14 ± 6.97 |  |  |  | 22.50 ± 8.09 |  |  |  |
| Comorbidities (n, %) | 23, 27.38% | — |  |  | 6, 42.86% | — |  |  |

NOTE. ADHD, attention-deficit/hyperactivity disorder; HC, healthy controls; SD, standard deviation; FSIQ, full-scaled intelligence quotient.

### MRI Acquisition and Preprocessing

All MRI data were acquired on a 3.0-Tesla MR system (General Electric; Discovery MR750) in the Center for Neuroimaging in Peking University Sixth Hospital. The 8-min resting-state fMRI data were acquired using an Echo planar Imaging (EPI) sequence with the following parameters: 43 axial slices, slice thickness =3.2 mm, slice skip= 0 mm, repetition time (TR) = 2000 ms, echo time (TE) =30 ms, flip angle (FA) = 90°,matrix=64×64, field of view (FOV) = 220×220 mm^2^, 240 volumes. High resolution T1-weighted anatomical images were acquired with the following parameters: 180 sagittal slices, slice thickness = 1 mm, slice skip= 0 mm, TR = 6.66 ms, TE = 2.93 ms, TI = 450 ms, FA= 8°, FOV = 256×256 mm^2^, matrix = 256×256.

The BOLD fMRI images were preprocessed using Statistical Parametric Mapping (SPM8, http://www.fil.ion.ucl.ac.uk/spm) and Data Processing Assistant for Resting-State fMRI [DPARSF, (Yan and Zang 2010)]. Briefly, we first discarded the first 10 volumes of each participant, corrected the acquisition time delay through slice timing and head motion through realignment to the first volume. No subject was excluded under a head motion criterion of 3 mm and 3°. The individual T1-weighted images were coregistered to the mean functional image after motion correction using a linear transformation (Collignon et al. 1995) and were then segmented into gray matter (GM), white matter, and cerebrospinal fluid tissue maps with SPM’s a priori tissue maps as reference by using a unified segmentation algorithm (Ashburner and Friston 2005). The resultant GM, white matter, and cerebrospinal fluid images were further nonlinearly registered into the Montreal Neurological Institute (MNI) space with the information estimated in unified segmentation and then averaged across all subjects to create custom GM, white matter, and cerebrospinal fluid templates. A customer-generated grey matter mask were then generated through thresholding grey matter probability > 25%. We then applied the transformation parameters estimated during unified segmentation to the motion-corrected functional volumes and resampled the transformational functional images to 3-mm isotropic voxels that are the minimum spatial resolution. The normalized functional images further underwent spatial smoothing with a 4-mm full width at half maximum (FWHM) Gaussian kernel and removal of linear trends. Temporal band-pass filtering (0.01–0.1Hz) was performed on the time series of each voxel. Several nuisance variables, including Friston’s 24 head motion parameters (Friston et al. 1996), the averaged signal from white matter, cerebrospinal fluid tissue and global signal, were further removed through multiple linear regression analysis to reduce the effects of nonneuronal signals.

### Connectome-wide Association Analysis

After preprocessing, the baseline resting-state fMRI data (N_ADHD_ =84, N_control_=89) were analyzed with a whole-brain voxel-based analysis approach (Brady et al., 2019; Shehzad et al., 2014) to get the voxel-based biomarker (*i*.*e*. ADHD vs control) for adults with ADHD. Specifically, for each subject, we calculated the voxel-based *v*×*v* correlation matrix (*v* = 53, 970) through temporal Pearson correlation of blood oxygen level dependent signals between each pair of grey matter voxles within the customer-generated grey matter mask. Then, the individual differences for each group in functional connectivity were calculated through a metric of distance, which were calculated as 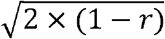, where *r* is the Pearson correlation of the connectivity pattern of each voxel within the mask (i.e., each voxel’s correlation with the rest of the brain) for every possible pairing of participants. In this way, we get an *n*×*n* connectivity distance matrix for each voxel, where *n* is the number of participants within the same group. The third step used diagnosis (*i*.*e*. ADHD=1, control=0) for each participant combined with the connectivity distance matrix to produce a pseudo-F statistic through MDMR, which characterizes the grouping ability to describe the similarity of the functional connectivity. The standard permutation flow of 15, 000 times was then employed the get the significance of the pseudo-F statistic in which the study subjects’ labels was randomly generated for each time to generated the null distribution. The pseudo-F statistic from the original data was then referred to this simulated distribution to obtain a p-value. The threshold (voxel-wise: *P*_permutation_<0.001; cluster size >50 voxels) was used for multiple comparisons correction.

After the identification of the significant clusters through the MDMR methods, a complimentary seed-based functional connectivity analysis was employed to explore detailed information of the connectional changes. We defined two seed regions of interest (ROIs) of all voxels within the regional clusters showing significant group-differences on connectivity similarity between groups, the precuneus and left middle temporal cluster, which was defined as the seed regions. For each region, we performed individual functional connectivity analysis by correlating the mean time series of the seed ROI with those of all voxels within the gray matter mask. A Fisher’s r-to-z transformation was further applied to improve the normality of the resulting correlation coefficient. Two-sample *t*-test (i.e., ADHD vs HC) was conducted to explore the group differences. The significance threshold (voxel-wise: *P* < 0.001; cluster-corrected: *P* < 0.05) was employed in this step. Notably, we employed a predefined subnetwork template which was adapted from Yeo et al. (2015) with 7 subnetworks to grouping the voxels showed disrupted functional connections with the seed ROIs. For each seed ROI, we calculated the percentages of identified voxels within every subnetwork.

### Correlation analysis between changes of clinical performances and neuroimaging biomarkers after CBT treatment

After the identification of brain connectivity-based biomarkers, we also explored the changes of those connections with clinical performances for the treatment effects of cognitive behavioral therapy (CBT). The data of 14 adults with ADHD with available information of both baseline and follow-up were included in the analysis. We calculated the association between between the changes of ADHD total scores before and after CBT treatment (*i*.*e*. Score_follow-up_ – Score_baseline_) and the changes of the 21 identified biomarkers (*i*.*e*. FC_follow-up_ – FC_baseline_) across subjects through Pearson correlation. Changes of ADHD inattention scores and hyperactivity/impulsivity scores were also calculated. Notably, we also collected the follow-up brain imaging data of 25 HCs after 12 weeks to excluding the possible influences of test-retest reliability.

### Analysis of MAOA and MAOB Genotype

We genotyped two monoamine oxidase genes, *MAOA* and *MAOB*. The monoamine oxidase A *(MAOA)* was known as the ADHD distinguishable genotype (Nymberg et al., 2013). *MAOB* has been suggested to be more close with adults with ADHD (Bonvicini et al., 2018). Four SNPs (*i*.*e*.rs1465108, rs5906883, rs6323, rs5905859) of *MAOA* and eight SNPs (*i*.*e*. rs1799836, rs10521432, rs2239449, rs2283729, rs6651806, rs2283727, rs3027441, rs5952671) of *MAOB* were genotyped using the Sequenom MassARRAY^®^ platform (Sequenom, San Diego, CA, USA).

We coded the SNPs as following: (1) heterozygote coded as ‘1’ (*e*.*g*. ‘AC’ coded as ‘1’); (2) homozygous coded as ‘2’ or ‘0’ which follows the letter’s order (*e*.*g*. ‘AA’ & ‘CC’ coded as ‘2’for ‘AA’; ‘0’for ‘CC’). We investigated whether *MAOA* and *MAOB* genotype could be used for stratification by testing the association between the two genetic genotypes and the identified biomarkers. The function of “partialcorr” in Matlab was employed to investigate the partial *Pearson* correlation coefficient through the gender respective (i.e., male and female) analysis with the mean FD, age, FSIQ, education year and ADHD diagnosis (i.e., ADHD=1, Control=0) as the covariates.

## Results

### Network Discovery With Multivariate Distance Matrix Regression (MDMR)

MDMR reveals the differences between functional connectivity patterns between adult ADHD and healthy control (N_ADHD_ =84, N_control_=89) were concentrated in the precuneus cluster (number of voxels: 141, Peak MNI coordinate: 6, −51, 3; including the left precuneus, right precuneus, cingulate gyrus and posterior cingulate; **Figure 1.A**) and the left middle temporal cluster (number of voxels: 72, Peak MNI coordinate: −60, −57, −6; including the temporal lobe,middle temporal gyrus, superior temporal gyrus; **Figure 1.B**).

**Figure 1.**
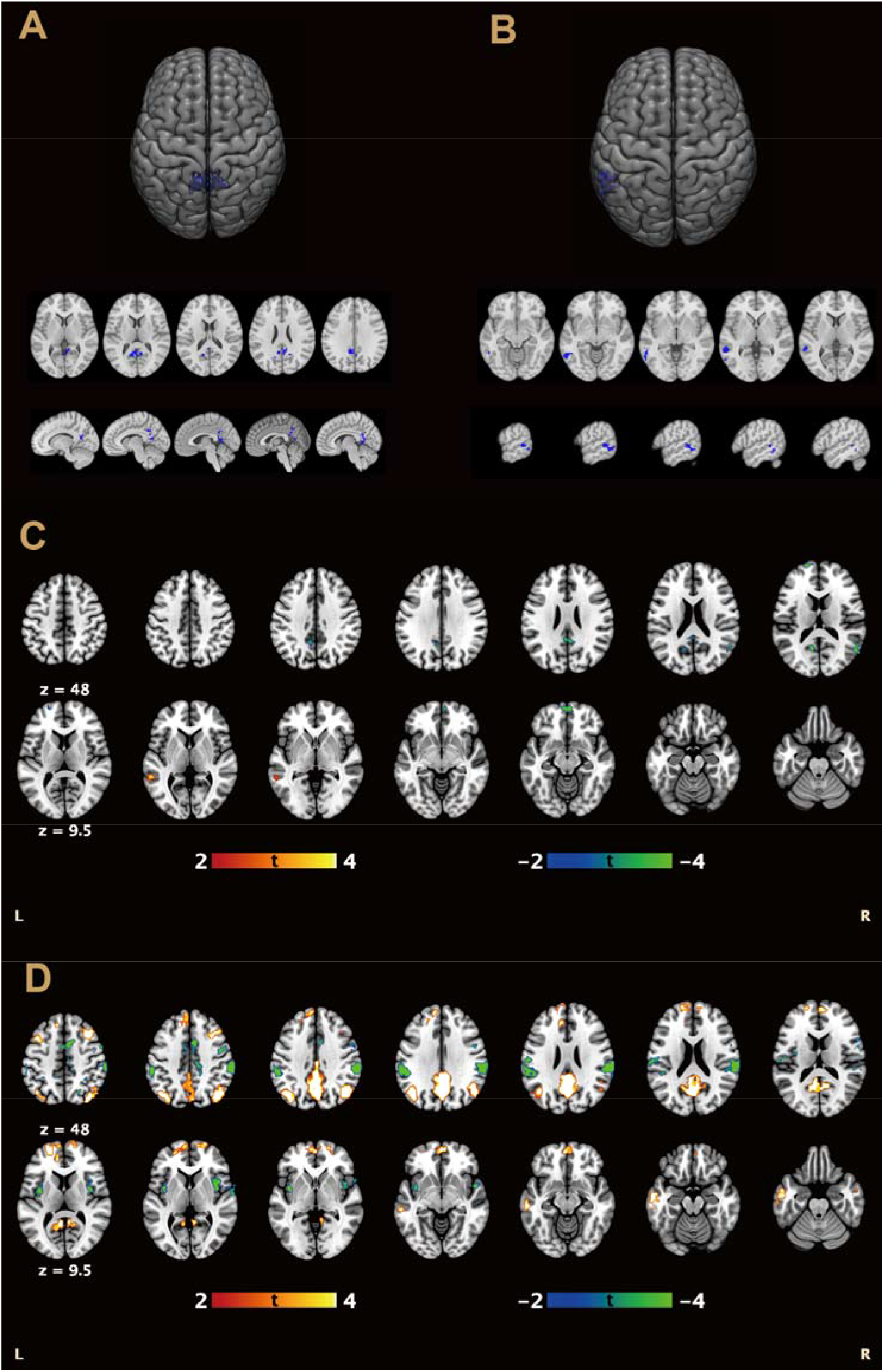
Neural basis of adults with ADHD. **(A)** Seed ROI: Left precuneus cluster; **(B)** Seed ROI: Left middle temporal cluster; **(C-D)** Post whole-brain t-value map (Precuneus cluster based for C, Left middle temporal cluster based for D);

After that, the post hoc seed-based connectivity analysis found 6 precuneus-based FCs and 15 temporal-based FCs showed significant differences between ADHD and HC groups, and these functional connectivity were considered as the biomarkers distinguish between adult ADHD and healthy control (voxel-wise: *P*_two-tail_<0.001; cluster-corrected: *P*_two-tail_<0.05; **Table 2**; **Figure 1.C, D**). Specifically, for both the ADHD and HC groups, the precuneus ROI showed positive connectivity with PCC/precuneus, left calcarine cortex, medial frontal gyrus and left superior frontal gyrus. However, the strength of these correlations was significantly lower in the ADHD than the HC group. Based on the subnetwork template of Yeo, most functional connection disruptions were located within the DMN (78.9%). For left middle temporal ROI, the changed connections could be summed in two groups. One is these linking with default mode network regions, including angular, middle frontal gyrus, precuneus and inferior temporal gyrus, which were negative in HC group but positive in ADHD group. These functional connections were also mainly linked with the DMN (77.52%). The other ones is linking with dorsal attention network and ventral attention regions (29.29% and 48.36%, respectively), including insula, superior parietal gyrus, supramarginal gyrus, precentral gryus and SMA, which were positive in HC group but negative in ADHD group. See **Table 3** and **Figure 2. A-C** for details.

**Table 2.**
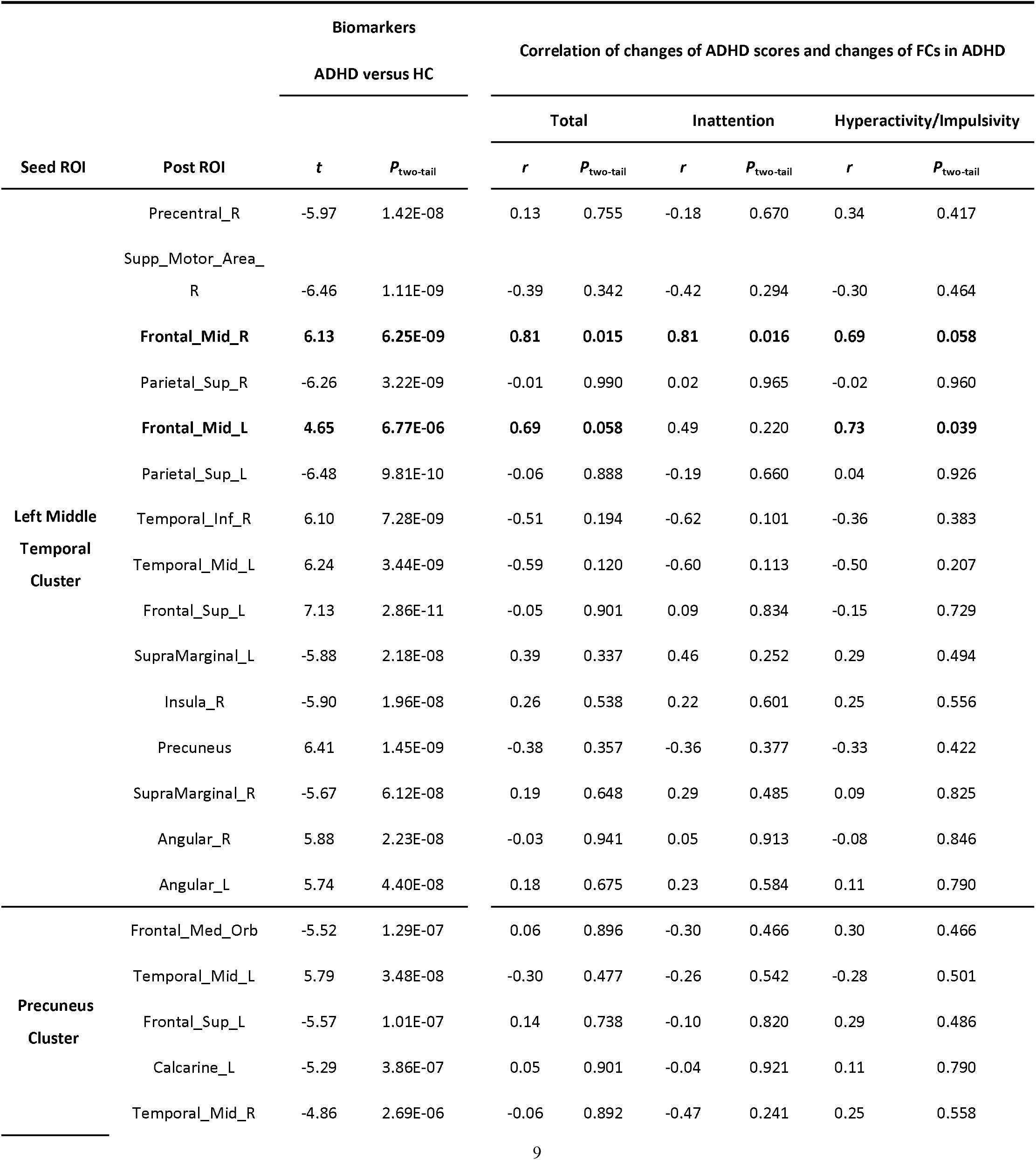
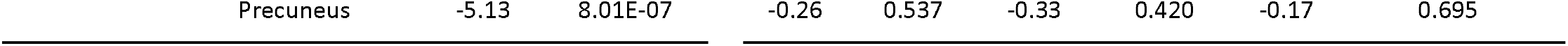
Biomarkers of adults with ADHD and the correlation between Change of ADHD scores and Change of FCs in ADHD group.

**Table 3.** Grouping of the voxels with significant altered seed-based FCs between ADHD and healthy controls according to the Yeo template.

|  |  | Visual Network | Somatomotor Network | Dorsal Attention Network | Ventral Attention Network | Limbic Network | Frontoparietal Network | Default Mode Network |
| --- | --- | --- | --- | --- | --- | --- | --- | --- |
| Temporal-based increased clusters | Number of voxels | 132 | 0 | 28 | 5 | 45 | 300 | 1759 |
|  | Percentage | 5.82% | 0% | 1.23% | 0.22% | 1.98% | 13.22% | **77.52%** |
| Temporal-based decreased clusters | Number of voxels | 7 | 309 | 490 | 809 | 0 | 56 | 2 |
|  | Percentage | 0.42% | 18.47% | **29.29%** | **48.36%** | 0% | 3.35% | 0.12% |
| Precuneus-based clusters | Number of voxels | 5 | 1 | 21 | 0 | 18 | 1 | 172 |
|  | Percentage | 2.29% | 0.46% | 9.63% | 0% | 8.26% | 0.46% | **78.9%** |

**Figure 2.**
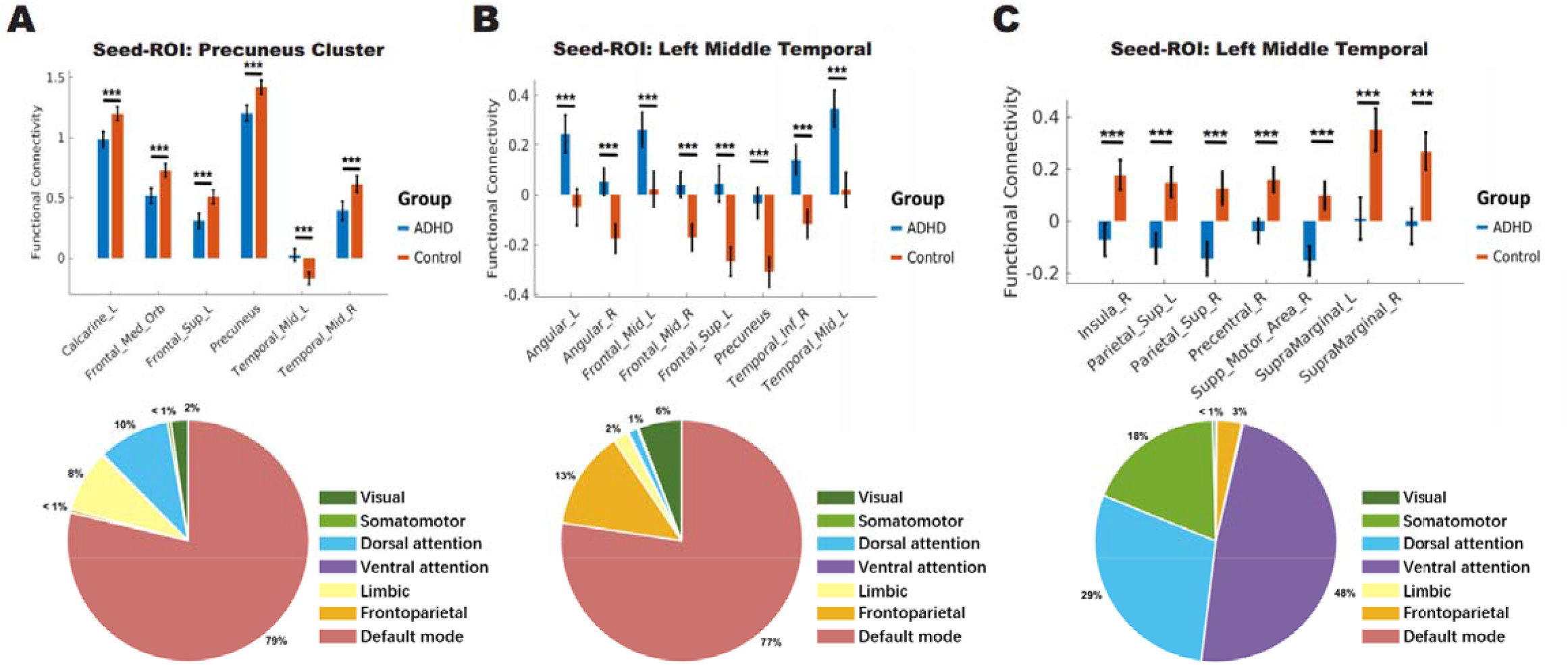
Significant changes of seed-based FCs between ADHD and healthy controls. **(A)** Precuneus cluster based FCs; **(B-C)** Left middle temporal cluster based FCs (increased for B; decreased for C); * *P* < 0.05, ** *P* < 0.01, *** *P* < 0.001.

### Effects of CBT treatment on identified biomarkers

The correlation analysis between the clinical performances (*i*.*e*. ADHD total Score_follow-up_ – Score_baseline_) and change of identified biomarkers (*i*.*e*. FC_follow-up_ – FC_baseline_) after treatment found the FC between the left middle temporal and right middle frontal cluster significant correlated to the change of ADHD scores (*r*=0.81, *P*_two-tail_ =0.015 for Total, **Figure 3.A**; *r*=0.81, *P*_two-tail_ = 0.016 for Inattention; **Table 2**). The FC between the left middle temporal and left middle frontal cluster marginally significant correlated to the change of ADHD scores (*r*=0.69, *P*_two-tail_ = 0.058 for Total, **Figure 3.B**; *r* = 0.73, *P*_two-tail_ = 0.039 for Hyperactivity/Impulsivity; **Table 2**)

**Figure 3.**
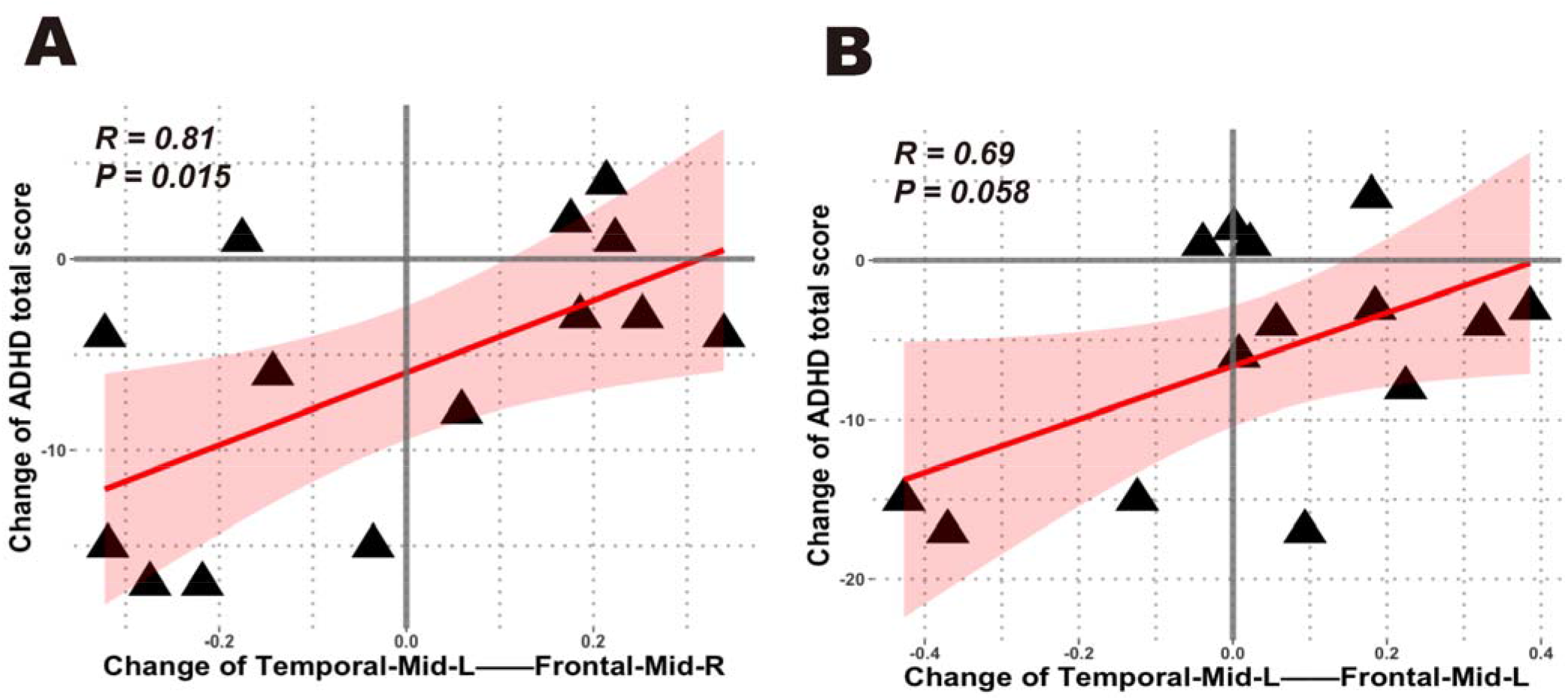
Significant correlations between the changes of ADHD total score and FCs before and after CBT treatment.

As a comparison, the two-sample *t*-test was employed to test the difference of healthy control from baseline to follow-up (*i*.*e*. baseline vs follow-up; N=25). 20 of 21 identified FCs showed no significant difference between baseline and follow-up time points (i.e., *P*_two-tail_> 0.05) in healthy control. Only the FC between precuneus cluster and right middle temporal has mild significant difference (*t*=2.15, *P*_two-tail_ =0.037) between baseline and follow-up. This findings indicating the changes in ADHD before and after CBT were not caused by the time effect.

### Analysis of MAOA and MAOB Genotype

We investigated whether *MAOA* and *MAOB* genotype might be used for testing the association between the two genetic genotypes and the 21 identified biomarkers.

For male (N_male_=107, **Figure 4.A**), the FC between precuneus cluster and left calcarine correlated to the *MAOB* genotype (*r*=−0.27, *P*_two-tail_=0.007 for rs2239449; *r*=0.20, *P*_two-tail_=0.041 for rs2283727; *r*=0.27, *P*_two-tail_=0.007 for rs3027441; *r*=−0.27, *P*_two-tail_=0.007 for rs5952671). The FC between precuneus cluster and right middle temporal gyrus significant correlated to the *MAOA* genotype (*r*=−0.22, *P*_two-tail_=0.030 for rs1465108; *r*=−0.21, *P*_two-tail_=0.037 for rs5906883; *r*=−0.22, *P*_two-tail_=0.027 for rs6323; *r*=0.23, *P*_two-tail_=0.023 for rs5905859) and *MAOB* genotype (*r*=−0.26, *P*_two-tail_= 0.009 for rs10521432; *r*= −0.22, *P*_two-tail_= 0.027 for rs2239449; *r*=−0.25, *P*_two-tail_=0.010 for rs2283729; *r*=0.25, *P*_two-tail_=0.010 for rs6651806; *r*=0.30, *P*_two-tail_=0.002 for rs2283727; *r*=0.22, *P*_two-tail_=0.027 for rs3027441; *r*=−0.22, *P*_two-tail_=0.027 for rs5952671). The FC between left middle temporal cluster and right inferior temporal gyrus significant correlated to the *MAOB* genotype (*r*=0.28, *P*_two-tail_=0.005 for rs1799836; *r*=0.20, *P*_two-tail_=0.045 for rs2283729; *r*=−0.20, *P*_two-tail_=0.045 for rs6651806). The FC between left middle temporal cluster and dorsolateral of left superior frontal gyrus, significant correlated to the *MAOB* genotype (*r*=0.20, *P*_two-tail_=0.045 for rs10521432; *r*=0.20, *P*_two-tail_= 0.043 for rs2239449; *r*=−0.20, *P*_two-tail_= 0.043 for rs3027441; *r*=0.20, *P*_two-tail_= 0.043 for rs5952671).

**Figure 4.**
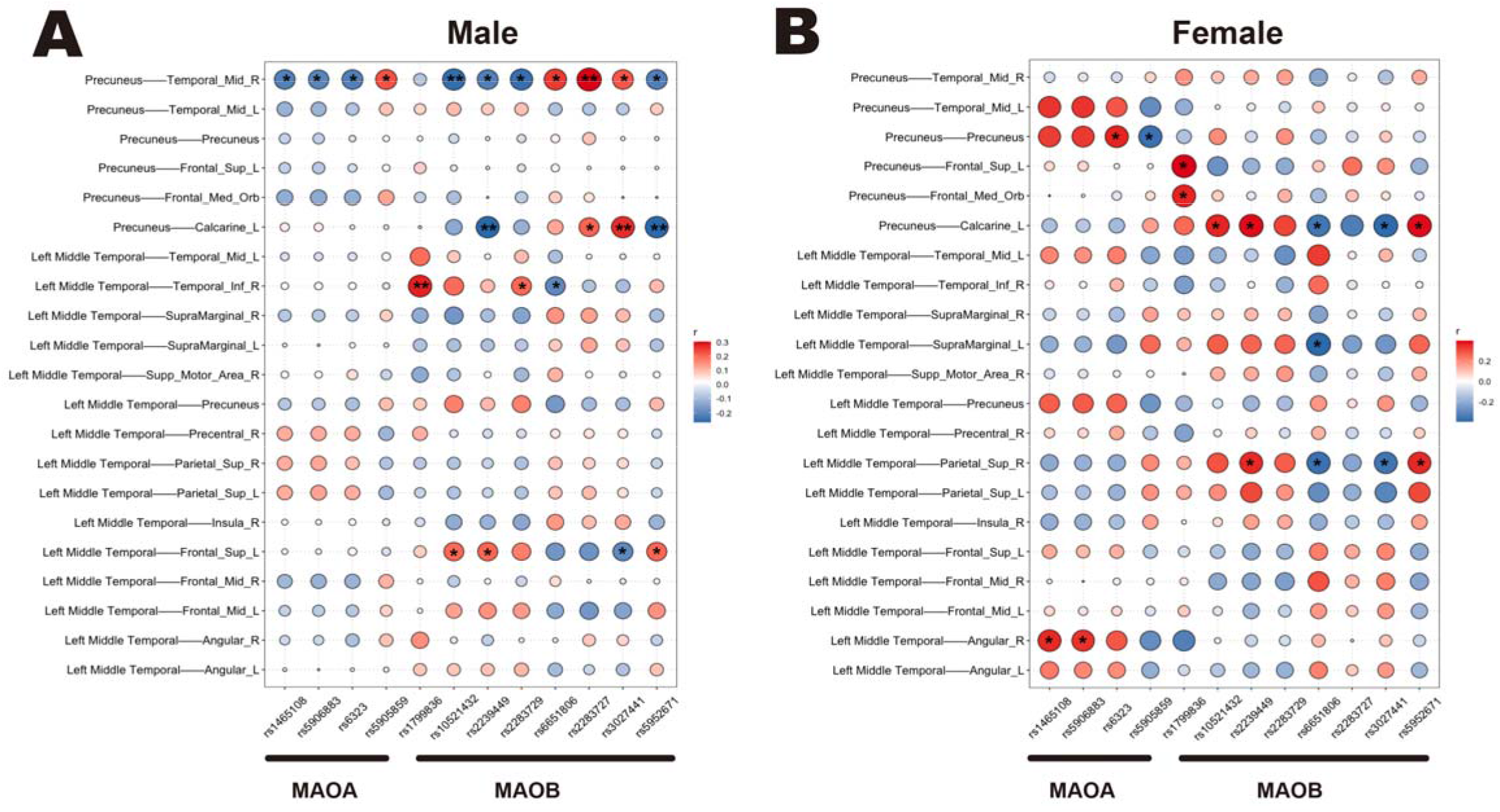
Correlation between MAO genotypes and identified baseline-FCs with significant group differences by gender respectively (male for A; female for B). * *P*< 0.05, ** *P*< 0.01, *** *P* < 0.001.

For female (N_female_=38, **Figure 4.B**), the FC between precuneus cluster and left calcarine significant correlated to the *MAOB* genotype (*r*=0.37, *P*_two-tail_=0.036 for rs10521432; *r*=0.39, *P*_two-tail_=0.025 for rs2239449; *r*=−0.36, *P*_two-tail_=0.042 for rs6651806; *r*=−0.39, *P*_two-tail_=0.025 for rs3027441; *r*=0.39, *P*_two-tail_=0.025 for rs5952671). The FC between left middle temporal cluster and right superior parietal gyrus significant correlated to the *MAOB* genotype (*r*=0.36, *P*_two-tail_=0.041 for rs2239449; *r*=−0.37, *P*_two-tail_=0.035 for rs6651806; *r*=−0.36, *P*_two-tail_= 0.041 for rs3027441; *r*=0.36, *P*_two-tail_=0.041 for rs5952671). The FC between precuneus cluster and precuneus significant correlated to the *MAOA* genotype (*r*=0.37, *P*_two-tail_=0.036 for rs6323; *r*=−0.37, *P*_two-tail_=0.036 for rs5905859). The FC between left middle temporal cluster and right angular significant correlated to the *MAOA* genotype (*r*=0.36, *P*_two-tail_= 0.044 for rs1465108; *r*=0.35, *P*_two-tail_=0.048 for rs5906883).

The more information about the correlation between the 21 identified FCs and two genotypes could be found in **Figure 4** for detail.

## Discussion

Our present study performed data-driven voxel-based analyses to identify connectome-wide significant brain functional features for adults with ADHD, an extreme-phenotype of ADHD. Remarkably, two clusters indicated alteration of functional connectivity in ADHD patients, including the left precuneus cluster and the left middle temporal cluster. The subsequent post hoc whole-brain voxel-based connectivity map with those two clusters as the new seeds indicated 21 FCs as potential neural biomarkers for adults with ADHD. After CBT, two of these ADHD-altered FCs could be ameliorated accompanied with improvement of ADHD core symptoms, indicating the good predictive validity of these brain functional features. In addition, imaging genetic analyses indicated close relationship between the observed brain functional alterations and ADHD-risk genes, indicating the effective conduct validity.

The left precuneus cluster identified using MDMR included the left precuneus, right precuneus, cingulate gyrus and posterior cingulate (PCC). Precuneus/PCC was involved in the hub brain regions of DMN as one part of its posterior components. In our present study, the precuneus seed-based analyses indicated decreased positive connectivity with other components of DMN in ADHD compared to HC. This result was consistent with previous reports of atypical default-mode connectivities in ADHD (Castellanos et al., 2008; Fair et al., 2010), indicating the altered FCs between posterior and anterior default-mode components. In another study by Uddin et al. (2008), the network homogeneity analyses also showed decreased integrity of DMN in ADHD, especially that between precuneus and other components of DMN. These consistent evidence supported the complex and significant role of precuneus/PCC in the integration between anterior and posterior default-mode components. The altered homogeneity and integration of DMN in ADHD subjects might influence the switch from DMN to cognitive networks, that further leads to deficits in cognitive performance, such as attention, working memory, inhibition control and so on. Interestingly, this decreased intra-DMN connectivity in adults with ADHD could be improved by methylphenidate (Picon et al., 2020). In addition, methyphenidate could normalize the abnormal patterns of task-related DMN deactivation in ADHD (Liddle et al., 2011). However, as indicated in our present study, CBT did not improve the abnormal pattern of intra-DMN connectivity, wheras genetic association was indicated with ADHD-related genes. It is worth to explore whether the intra-DMN dysfunction is more sensitive to medication than CBT. However, we should note that the sample size included for CBT-image analyses is small, and the results should be interpreted with caution and need to be validated. In addition to functional impairments, significant alteration in multiple structural indices of precuneus and PCC has been indicated in prior findings of both children and adults with ADHD, including decreased volume, cortical thickness and surface area (Firouzabadi et al., 2021). It is also worthy of further exploration to elucidate whether and how these structural and functional abnormalities in DMN regions jointly participated in the pathogenesis of ADHD.

Another identified interesting region was the left middle temporal cluster including the temporal lobe, middle temporal gyrus and the superior temporal gyrus. Further seed-based connectivity analysis indicated different and even opposite direction signals between adults with ADHD and HC. In detailed, the FCs between left middle temporal and default-mode regions, such as precuneus, angular gyrus, middle frontal cortex, were negative in healthy controls, wheras these FCs were almost positive in adults with ADHD. Similarly, different from the positive FCs in HC group between left middle temporal and regions of attention networks including insula, superior parietal lobe, adults with ADHD showed weak negative FCs. In the existing literature, the negative relationship between DMN and control regions is intrinsically represent in the human brain (Weissman et al., 2006). Our present study supported this negative correlation in both ADHD and HCs to some extent. As discussed in Rubia (2018), both hyper-engaged DMN and hypo-engaged task-relevant networks contributed to cognitive impairment. Correspondingly, atypical inter-network connectivity between DMN and cognitive networks has been illustrated in ADHD patients, which was closely associated impairments in multiple cognitive performance including attention and response control (Duffy et al., 2021). Different from the direct calculation of inter-connectivity between DMN and task-relevant networks, our present study indicated middle temporal gyrus as the key “bridge” region linked DMN and attention networks. Notably, the FCs changed between the left temporal and right/left middle frontal cluster were positively correlated with the improvement of ADHD core symptoms after CBT, especially the inattention symptom. This suggested that CBT might improve the ADHD symptoms by influencing the functional connectivity between temporal lobe and default-mode regions. In our previous study, the regional homogeneity (ReHo) values of parahippocampal cluster (including middle temporal gyrus) in adults with ADHD increased after CBT (Cao et al., 2017). These results supported the importance of brain functional alteration with middle temporal gyrus in the brain mechanisms of ADHD from the predictive validity. Meanwhile, genetic analyses also supported the involvement of middle temporal-related functional impairment in ADHD from the constructive validity.

Middle temporal gyrus (MTG) has been generally suggested to play important role in visual processing. Nevertheless, it is also involved in multiple brain networks, including DMN and attention networks (Lin and Gao, 2016). One recent EEG study indicated increased temporal lobe beta activity in boys with ADHD and increased connectivity between the left middle frontal gyrus and fusiform gyrus of the temporal lobe, while Beta activity represents attention level (Chiang et al., 2020). A recent task-based fMRI study showed main effects of middle temporal gyrus/superior temporal gyrus on reading and attentional control (Arrington et al., 2019). As illustrated in the study of Hoogman et al. (2019), lower surface area values and cortical thickness of multiple brain regions were found in children with ADHD, including temporal regions. Further, the decreased surface area of middle temporal gyrus showed significant association with inattention symptoms. Although these case-control brain structural difference was nonsignificant in adults, the delayed structural maturation during childhood might influence the construction and dynamic development of functional connectivity of temporal gyrus with other brain regions. To further explore and validate the key role of middle temporal gyrus in the relationship between DMN and cognitive task-relevant networks, especially attention networks, fusion analysis of both resting-state and task-based FCs could help to elucidate more comprehensively. In addition, integration of state and dynamic FCs analyses should also be considered.

In summary, our present study used multivariate distance matrix regression (MDMR) to conduct a whole-brain voxel-based analysis and explore the potential imaging biomarkers of adults with ADHD. Further treatment effects and genetic analyses validated the validity of these findings from predictive and constructive perspectives. However, some limitations should be considered. First, we did not explore the potential gender difference in our present analyses. Instead, ADHD and HCs groups were gender-matched to control the potential confounding influence of gender. Second, the follow-up subjects were with a small sample size, which was only a secondary analyses. A rigorous design of treatment-imaging study could be considered in the future to validate our primary findings. Third, for the imaging genetic studies, we only analyzed limited genetic variants of two ADHD-related genes. More genetic variants analyzed and different genetic parameters (i.e. polygenic risk score) should be considered. Finally, multimodal imaging features could be further combined for analyses to establish a more comprehensive brain alterations in adults with ADHD.

## Conclusion

Our hypothesis-free MDMR analyses indicated the atypical patterns of brain functional connectivity in adults with ADHD, involving both within-DMN and DMN-cognitive networks. More significantly, middle temporal gyrus might be a key ‘bridge’ region which linked DMN and task-relevant networks, which could be an important biomarker for both ADHD pathogenesis and treatment effects. The more comprehensive exploration of middle temporal gyrus is needed and of great significance.

## Author Contributions

Lu Liu, Fang Huang, Qiujin Qian and Miao Cao contributed conception and design of the study; Di Chen, Miao Cao, Tianye Jia, Xuan Bu and Fang Huang performed the statistical analysis. Lu Liu, Di Chen, Fang Huang, Mengjie Zhao, Meirong Pan, Miao Cao, Qiujin Qian and Yufeng Wang interpreted the results and wrote the manuscript. Jianfeng Feng revised the manuscript critically. All authors contributed to manuscript revision, read and approved the submitted version.

## References

Bonvicini C, Faraone SV, Scassellati C. Common and specific genes and peripheral biomarkers in children and adults with attention-deficit/hyperactivity disorder. World J Biol Psychiatry. 2018 Mar;19(2):80–100.

Brady RO Jr, Gonsalvez I, Lee I, Öngür D, Seidman LJ, Schmahmann JD, Eack SM, Keshavan MS, Pascual-Leone A, Halko MA. Cerebellar-Prefrontal Network Connectivity and Negative Symptoms in Schizophrenia. Am J Psychiatry. 2019 Jul 1;176(7):512–520.

Castellanos FX, Margulies DS, Kelly C, Uddin LQ, Ghaffari M, Kirsch A,Shaw D, Shehzad Z, Di Martino A, Biswal B, Sonuga-Barke EJS, Rotrosen J, Adler LA and Milham MP. Cingulate-precuneus interactions:a new locus of dysfunction in adult attention-deficit/hyperactivitydisorder. Biological Psychiatry. 2008, 63: 332–337.

Chiang CT, Ouyang CS, Yang RC, Wu RC, Lin LC. Increased Temporal Lobe Beta Activity in Boys With Attention-Deficit Hyperactivity Disorder by LORETA Analysis. Front Behav Neurosci. 2020 Jun 30;14:85.

Cao QJ, Wang XL, Qu S, Wang P, Wu ZM, Sun L, Wang YF. Effects of cognitive-behavioral therapy on regional homogeneity changes in adults with attention-deficit/hyperactivity disorder. Chinese Mental Health Journal. 2016, 30:

Chen Q, Brikell I, Lichtenstein P, Serlachius E, Kuja-Halkola R, Sandin S, Larsson H. Familial aggregation of attention-deficit/hyperactivity disorder. J Child Psychol Psychiatry. 2017 Mar;58(3):231–239.

Cortese S, Aoki YY, Itahashi T, Castellanos FX, Eickhoff SB. Systematic Review and Meta-analysis: Resting-State Functional Magnetic Resonance Imaging Studies of Attention-Deficit/Hyperactivity Disorder. J Am Acad Child Adolesc Psychiatry. 2021 Jan;60(1):61–75.

Dobrosavljevic M, Solares C, Cortese S, Andershed H, Larsson H. Prevalence of attention-deficit/hyperactivity disorder in older adults: A systematic review and meta-analysis. Neurosci Biobehav Rev. 2020 Nov;118:282–289.

Duffy KA, Rosch KS, Nebel MB, Seymour KE, Lindquist MA, Pekar JJ, Mostofsky SH, Cohen JR. Increased integration between default mode and task-relevant networks in children with ADHD is associated with impaired response control. Dev Cogn Neurosci. 2021, 50: 100980.

Fair DA, Posner J, Nagel BJ, Bathula D, Dias TG, Mills KL, Blythe MS, Giwa A, Schmitt CF, Nigg JT. Atypical default network connectivity in youth with attention-deficit/hyperactivity disorder. Biol Psychiatry. 2010, 68:1084–1091.

Firouzabadi FD, Ramezanpour S, Firouzabadi MD, Yousem IJ, Puts NAJ, Yousem DM. Neuroimaging in Attention-Deficit/Hyperactivity Disorder: Recent Advances. AJR Am J Roentgenol. 2021 Aug 18. doi: 10.2214/AJR.21.26316.

Franke B, Faraone SV, Asherson P, Buitelaar J, Bau CH, Ramos-Quiroga JA, Mick E, Grevet EH, Johansson S, Haavik J, Lesch KP, Cormand B, Reif A; International Multicentre persistent ADHD Collaboration. The genetics of attention deficit/hyperactivity disorder in adults, a review. Mol Psychiatry. 2012 Oct;17(10):960–87.

Gao Y, Shuai D, Bu X, Hu X, Tang S, Zhang L, Li H, Hu X, Lu L, Gong Q, Huang X. Impairments of large-scale functional networks in attention-deficit/hyperactivity disorder: a meta-analysis of resting-state functional connectivity. Psychol Med. 2019 Nov;49(15):2475–2485.

Huang F, Tang YL, Zhao M, Wang Y, Pan M, Wang Y, Qian Q. Cognitive-Behavioral Therapy for Adult ADHD: A Randomized Clinical Trial in China. J Atten Disord. 2019 Jul;23(9):1035–1046.

Lara C, Fayyad J, de Graaf R, Kessler RC, Aguilar-Gaxiola S, Angermeyer M, Demytteneare K, de Girolamo G, Haro JM, Jin R, Karam EG, Lépine JP, Mora ME, Ormel J, Posada-Villa J, Sampson N. Childhood predictors of adult attention-deficit/hyperactivity disorder: results from the World Health Organization World Mental Health Survey Initiative. Biol Psychiatry. 2009 Jan 1;65(1):46–54.

Liddle EB, Hollis C, Batty MJ, Groom MJ, Totman JJ, Liotti M. Task-related default mode network modulation and inhibitory control in ADHD: effects of motivation and methyphenidate. J Child Psychol Psychiatry. 2011, 52(7): 761–771.

Lin HY, Gau SS. Atomoxetine treatment strengthens an anti-correlated relationship between functional brain networks in medication-naïve adults with attention-deficit hyperactivity disorder: A randomized double-blind placebo-controlled clinical trial. Int J Neuropsychopharmacol. 2016;19(3):1–15.

McCarthy H, Skokauskas N, Mulligan A, Donohoe G, Mullins D, Kelly J, Johnson K, Fagan A, Gill M, Meaney J, Frodl T. Attention network hypoconnectivity with default and affective network hyperconnectivity in adults diagnosed with attention-deficit/hyperactivity disorder in childhood. JAMA Psychiatry. 2013 Dec;70(12):1329–37.

Nymberg C, Jia T, Lubbe S, Ruggeri B, Desrivieres S, Barker G, Büchel C, Fauth-Buehler M, Cattrell A, Conrod P, Flor H, Gallinat J, Garavan H, Heinz A, Ittermann B, Lawrence C, Mann K, Nees F, Salatino-Oliveira A, Paillère Martinot ML, Paus T, Rietschel M, Robbins T, Smolka M, Banaschewski T, Rubia K, Loth E, Schumann G; IMAGEN Consortium. Neural mechanisms of attention-deficit/hyperactivity disorder symptoms are stratified by MAOA genotype. Biol Psychiatry. 2013 Oct 15;74(8):607–14.

Pereira-Sanchez V, Franco AR, Vieira D, de Castro-Manglano P, Soutullo C, Milham MP, Castellanos FX. Systematic Review: Medication Effects on Brain Intrinsic Functional Connectivity in Patients With Attention-Deficit/Hyperactivity Disorder. J Am Acad Child Adolesc Psychiatry. 2020 Oct 27:S0890-8567(20)32054-2.

Picon FA, Sato JR, Anés M, et al. Methylphenidate alters functional connectivity of default mode network in drug-naive male adults with ADHD. J Atten Disord. 2020;24(3):447–455.

Rubia K. Cognitive neuroscience of attention-deficit hyperactivity disorder (ADHD) and its clinical translation. Front Hum Neurosci. 2018, 12:100.

Shehzad Z, Kelly C, Reiss PT, Cameron Craddock R, Emerson JW, McMahon K,Copland DA,Castellanos FX, Milham MP. A multivariate distance-based analytic framework for connectome-wide association studies. Neuroimage. 2014 Jun;93 Pt1(01):74–94.

Uddin LQ, Kelly AM, Biswal BB, et al. Network homogeneity reveals decreased integrity of default-mode network in ADHD. J Neurosci Methods, 2008,169(1):249–254.

Wang X, Cao Q, Wang J, Wu Z, Wang P, Sun L, Cai T, Wang Y. The effects of cognitive-behavioral therapy on intrinsic functional brain networks in adults with attention-deficit/hyperactivity disorder. Behav Res Ther. 2016 Jan;76:32–9.

Weissman DH, Roberts KC, Visscher KM, Woldorff MG. The neural bases of momentary lapses in attention. Nat Neurosci. 2006, 9: 971–978.

Yan CG, Zang YF. DPARSF: A MATLAB toolbox for “pipeline” data analysis of resting-state fMRI. Frontiers in Systems Neuroscience. 2010, 4: 13.

Yang Z, Kelly C, Castellanos FX, Leon T, Milham MP, Adler LA. Neural Correlates of Symptom Improvement Following Stimulant Treatment in Adults with Attention-Deficit/Hyperactivity Disorder. J Child Adolesc Psychopharmacol. 2016 Aug;26(6):527–36.

